# Systematic tissue-specific functional annotation of the human genome highlights immune-related DNA elements for late-onset Alzheimer’s disease

**DOI:** 10.1101/078865

**Authors:** Qiongshi Lu, Ryan L. Powles, Sarah Abdallah, Derek Ou, Qian Wang, Yiming Hu, Yisi Lu, Wei Liu, Boyang Li, Shubhabrata Mukherjee, Paul K. Crane, Hongyu Zhao

**Author notes:** These authors contributed equally to this work (Qiongshi Lu, Ryan L. Powles). To whom correspondence should be addressed: Dr. Hongyu Zhao, Department of Biostatistics Yale, School of Public Health 60 College Street, New Haven, CT, 06520, USA.

## Abstract

Continuing efforts from large international consortia have made genome-wide epigenomic and transcriptomic annotation data publicly available for a variety of cell and tissue types. However, synthesis of these datasets into effective summary metrics to characterize the functional non-coding genome remains a challenge. Here, we present GenoSkyline-Plus, an extension of our previous work through integration of an expanded set of epigenomic and transcriptomic annotations to produce high-resolution, single tissue annotations. After validating our annotations with a catalog of tissue-specific non-coding elements previously identified in the literature, we apply our method using data from 127 different cell and tissue types to present an atlas of heritability enrichment across 45 different GWAS traits. We show that broader organ system categories (e.g. immune system) increase statistical power in identifying biologically relevant tissue types for complex diseases while annotations of individual cell types (e.g. monocytes or B-cells) provide deeper insights into disease etiology. Additionally, we use our GenoSkyline-Plus annotations in an in-depth case study of late-onset Alzheimer’s disease (LOAD). Our analyses suggest a strong connection between LOAD heritability and genetic variants contained in regions of the genome functional in monocytes. Furthermore, we show that LOAD shares a similar localization of SNPs to monocyte-functional regions with Parkinson’s disease. Overall, we demonstrate that integrated genome annotations at the single tissue level provide a valuable tool for understanding the etiology of complex human diseases. Our GenoSkyline-Plus annotations are freely available at http://genocanyon.med.yale.edu/GenoSkyline.

**Author Summary:** After years of community efforts, many experimental and computational approaches have been developed and applied for functional annotation of the human genome, yet proper annotation still remains challenging, especially in non-coding regions. As complex disease research rapidly advances, increasing evidence suggests that non-coding regulatory DNA elements may be the primary regions harboring risk variants in human complex diseases. In this paper, we introduce GenoSkyline-Plus, a principled annotation framework to identify tissue and cell type-specific functional regions in the human genome through integration of diverse high-throughput epigenomic and transcriptomic data. Through validation of known non-coding tissue-specific regulatory regions, enrichment analyses on 45 complex traits, and an in-depth case study of neurodegenerative diseases, we demonstrate the ability of GenoSkyline-Plus to accurately identify tissue-specific functionality in the human genome and provide unbiased, genome-wide insights into the genetic basis of human complex diseases.

## Introduction

Large consortia such as ENCODE [1] and Epigenomics Roadmap Project [2] have generated a rich collection of high-throughput genomic and epigenomic data, providing unprecedented opportunities to delineate functional structures in the human genome. As complex disease research rapidly advances, evidence has emerged that disease-associated variants are enriched in regulatory DNA elements [3, 4]. Therefore, functional annotation of the non-coding genome is critical for understanding the genetic basis of human complex diseases. Unfortunately, categorizing the complex regulatory machinery of the genome requires integration of diverse types of annotation data as no single annotation captures all types of functional elements [5]. Recently, we have developed GenoSkyline [6], a principled framework to identify tissue-specific functional regions in the human genome through integrative analysis of various chromatin modifications. In this work, we introduce GenoSkyline-Plus, a comprehensive update of GenoSkyline that incorporates RNA sequencing and DNA methylation data into the framework and extends to 127 integrated annotation tracks covering a spectrum of human tissue and cell types.

To demonstrate the ability of GenoSkyline-Plus to systematically provide novel insights into complex disease etiology, we jointly analyzed summary statistics from 45 genome-wide association studies (GWAS; N_total_≈3.8M) and identified biologically relevant tissues for a broad spectrum of complex traits. We next performed an in-depth, annotation-driven investigation of Alzheimer’s disease (AD), a neurodegenerative disease characterized by deposition of amyloid-β (Aβ) plaques and neurofibrillary tangles in the brain. Late-onset AD (LOAD) includes patients with onset after 65 years of age and has a complex mode of inheritance [7]. Around 20 risk-associated genetic loci have been identified in LOAD GWAS [8]. However, our understanding of LOAD’s genetic architecture and disease etiology is still far from complete. Through integrative analysis of GWAS summary data and GenoSkyline-Plus annotations, we identified strong enrichment for LOAD associations in immune cell-related DNA elements, consistent with other data suggesting a crucial role for the immune system in AD etiology [9-11]. Jointly analyzing GWAS summary data for LOAD and Parkinson’s disease (PD), we identified substantial enrichment for pleiotropic associations in the monocyte functional genome. Our findings provide support for the critical involvement of the immune system in the etiology of neurodegenerative diseases, and suggest a previously unsuspected role for an immune-mediated pleiotropic effect between LOAD and PD.

## Results

### Identify tissue and cell type-specific functionality in the human genome

We use our previously established statistical framework to calculate the posterior probability of functionality for each nucleotide in the human genome [12]. Integrating tissue and cell-specific genomic functional data available through Epigenomics Roadmap Project [2], we make available GenoSkyline-Plus scores for 127 individual tissue annotation tracks (**Methods**; **S1 Table**). H3K4me3 and H3K9ac, known markers of open chromatin and active transcription [13], are shown to have the largest odds ratios of predicting functionality across the genome (**Fig 1A**). Identifying H3K4me3 and H3K9ac as strong indicators of genomic functionality is a finding consistent with previous studies of gene regulation through chromatin marks [14]. In contrast, H3K9me3, a well established repressive mark [13], has a reversed effect on genome functionality. The bimodal pattern of GenoSkyline scores [6] allows us to impose a score cutoff to robustly define the functional genome. Using a cutoff of 0.5, 3% of the genome is considered functional on average across all annotation tracks (**Fig 1B**). This functionality percentage varies from 1% in pancreatic islet cells to 8% in PMA-I stimulated T-helper cells. Our findings on functionality across all tracks are consistent with previous findings [12]; 34% of the intergenic human genome is predicted to be functional in at least one annotation track (**Fig 1C**). Additionally, coding regions of the genome are predicted to have much greater proportions of functionality in multiple tissues than intronic and intergenic regions.

**Fig 1.**
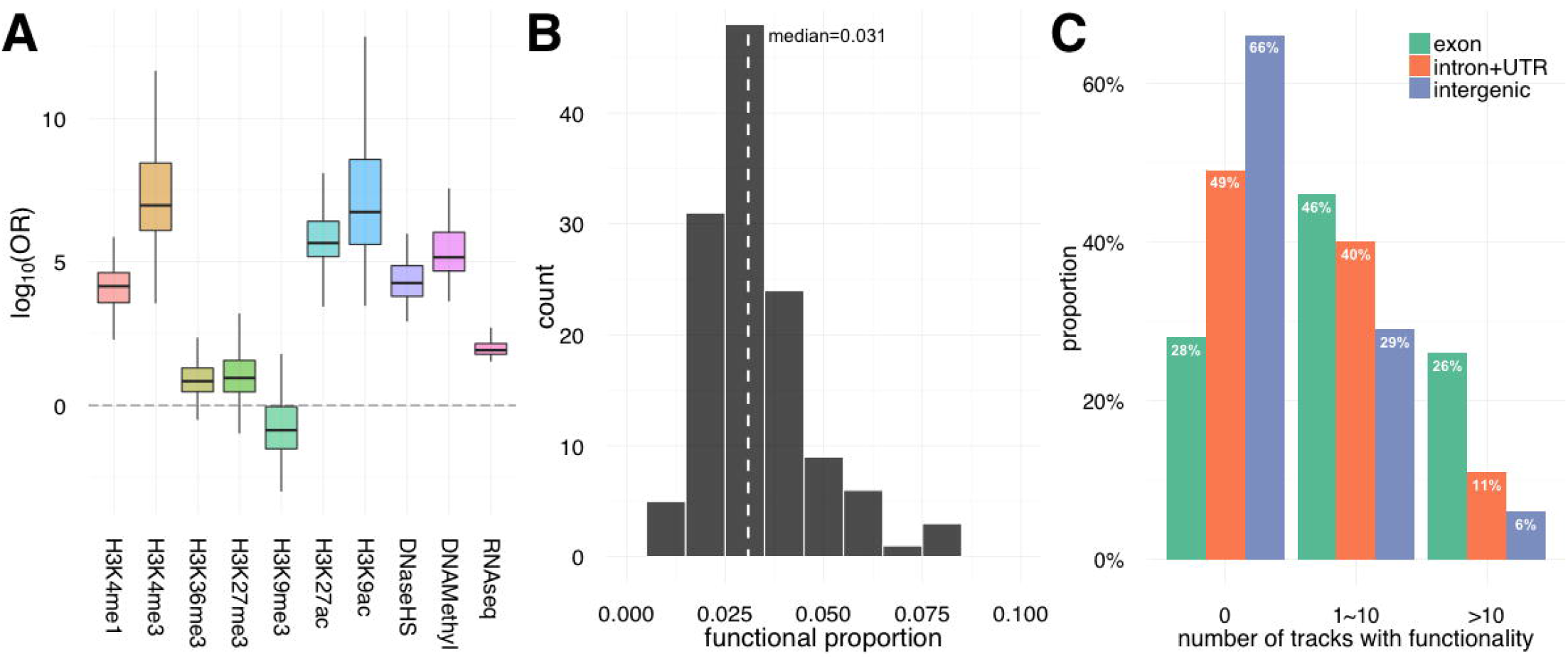
Basic characteristics of GenoSkyline-Plus annotation. (A) Odds ratio of predicting functionality. Each box represents the odds ratio for the same data type across 127 GenoSkyline-Plus tracks. (B) Histogram of predicted functional proportion across 127 annotation tracks. Dashed line marks the mean functional proportion. (C) Distribution of tracks with predicted functionality. For example, 26% of exon regions are predicted to be functional in more than 10 GenoSkyline-Plus tracks.

To assess the ability of GenoSkyline-Plus to capture tissue and cell-specific, non-coding functionality in the human genome, we consider a diverse set of known non-coding regulatory elements studied across the genome. To start, we examined microRNAs (miRNA), which are known to regulate a variety of cellular processes through the translational repression and degradation signaling of transcripts [15]. Recent work by Ludwig et al. profiled miRNA expression in 61 different human tissues and identified miRNAs with functionality unique to single tissues through a tissue specific index [16, 17] (TSI; **Methods**). We applied GenoSkyline-Plus scores to miRNA with tissue-specific functionality by calculating the total proportion of nucleotides predicted to be functional in each tissue. We next looked for which annotation tracks are able to predict the highest proportion of functionality for these known functional regions. The best predictors of high functionality for the three tissues with the largest sample sizes (i.e. brain, liver, and muscle) are tracks for brain structures, the liver track, and the muscle track, respectively (**Fig 2A**).

**Fig 2.**
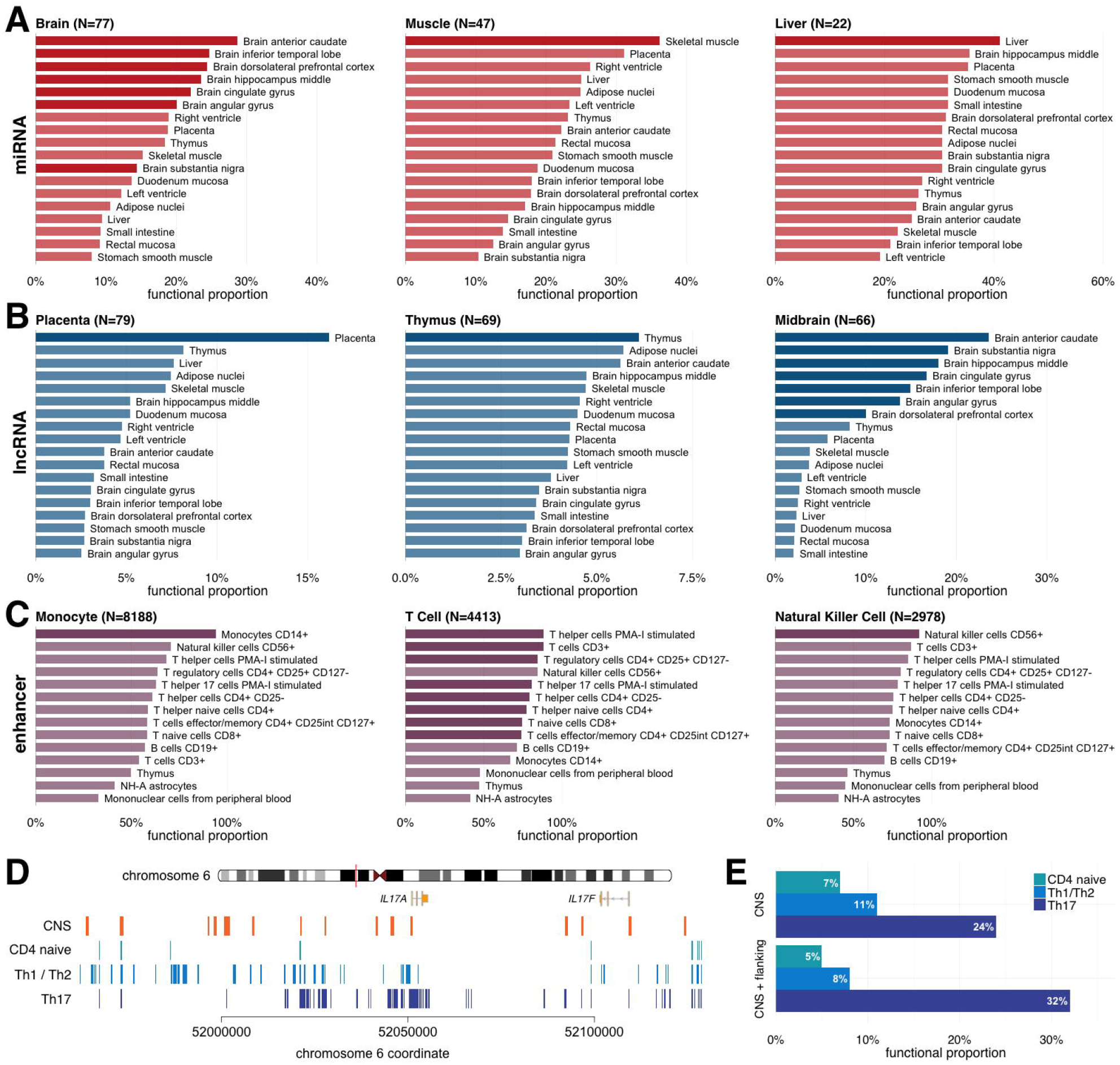
Identify tissue and cell type-specific functionality. Predicted functional proportions of different classes of previously identified tissue-specific non-coding elements. Sample sizes count the number of non-coding elements with specificity for the titled tissue or cell type (see **Methods**). Darker bars represent annotation tracks that physiologically match the tissue to which the corresponding set of non-coding elements are specific. **(A)** miRNAs with TSI > 0.75 identified in Ludwig et al. **(B)** lncRNAs with TSI > 0.75 identified in Derrien et al. **(C)** Enhancers with differential expression within a cell type facet identified by Andersson et al. **(D)** Predicted functional elements based on GenoSkyline-Plus annotations in the *IL-17A* LCR. Orange boxes mark identified CNS sites. **(E)** Predicted functional proportion in CNS sites and their 200-bp flanking regions across different T-cell subsets.

**Fig 3.**
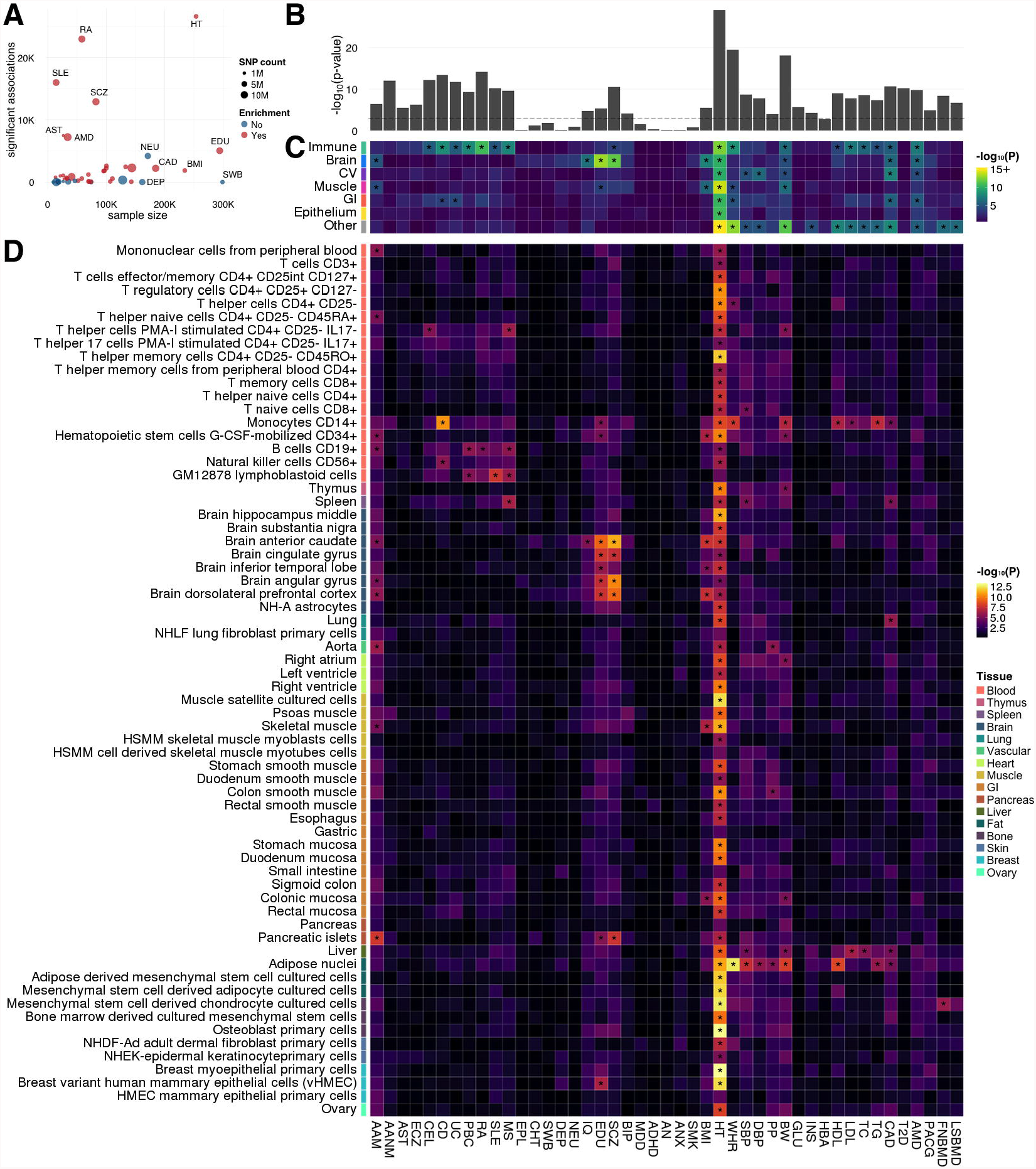
Enrichment analysis for 45 human complex traits. **(A)** Relationship between GWAS sample size, total count of significant associations, and signal enrichment in the functional genome. Traits significantly enriched in at least one annotation are highlighted in red. **(B)** Enrichment in the general functional genome predicted by GenoCanyon annotation. The dashed line marks the Bonferroni-corrected significance cutoff. **(C)** Enrichment across 7 broadly defined tissue tacks. Asterisks highlight significance after correcting for 45 traits and 7 tissues. **(D)** Enrichment in 66 tissue and cell tracks. Asterisks highlight significant enrichment after correcting for 45 traits and 66 annotations. Details for annotation tracks and different traits are summarized in **S2-3 Tables**.

We next examined long non-coding RNAs (lncRNA), another non-coding element known for its tissue-specific regulatory action [18]. Using a custom-designed microarray targeting GENCODE lncRNA, Derrien et al. profiled the activity of 9,747 lncRNA transcripts [19]. In order to reidentify and validate the set of lncRNA transcripts that are specific to their respective tissues, we calculated the previously described TSI and selected lncRNAs with expression specific to only a few cell types. Physiologically matching tracks show a higher proportion of predicted functionality than unmatched tracks in complex, heterogeneous tissue structures like the midbrain. More functionally uniform tissues, such as the thymus or placenta, show the highest functional proportion in matching annotation tracks (**Fig 2B**).

We also assessed enhancers, non-coding elements that can remotely regulate transcription of an associated promoter elsewhere on the genome with important roles in cell-type specificity [20]. We extracted tissue and cell type-specific enhancer facets identified through the FANTOM5 cap analysis of gene expression (CAGE) atlas and positive differential expression when compared against other defined facets [21]. To determine the utility of the large library of immune cells available in the Epigenomics Roadmap Project for which we developed annotation tracks, we focused on enhancer facets with differential CAGE expression in immune cells. While the method by which enhancers are defined to be differential in a facet is liberal (**Methods**) and does not imply facet-specific expression, GenoSkyline-Plus still showed outstanding ability to identify matching cell types. Indeed, matched annotation tracks for T-cells, natural killer cells, and monocytes show consistently higher functional proportions than other, non-matched immune cell annotation tracks (**Fig 2C**).

Finally, we present a case study of the *IL17A-IL17F* locus control region (LCR) in humans, a ∼200kb regulatory region surrounding the *IL17A* gene locus. *IL17A* encodes the primary secreted cytokine effector molecule IL-17 of T helper 17 (Th17) cells [22]. The LCR has been studied in mouse models and is found to contain many potential human-conserved intergenic regulatory elements that bind transcription factors that are essential for Th17 cell differentiation and effector function [23, 24]. Experimentally, these conserved noncoding sequences (CNS) acquire functionally permissive H3 acetylation marks at much greater magnitudes under Th17- inducing conditions than naïve or combined Th1 and Th2 populations [25]. Comparing annotation tracks for naïve CD4^+^ T-cells, differentiated Th17 cells, and differentiated Th1/Th2 cell populations, we identified highly Th17-specific functionality in the conserved regions of the human genome corresponding to known murine CNS regions (**Figs 2D and 2E**). CNS sites and their flanking regions showed substantially higher functional proportion in Th17 cells than in naïve CD4^+^ T-cells or Th1/Th2 cell subsets.

### Stratify heritability by tissue and cell type for 45 human complex traits

We jointly analyzed three tiers of annotation tracks that respectively represent the overall functional genome, 7 broad tissue clusters, and 66 tissue and cell types (**Methods**; **S2 Table**), with summary statistics from 45 GWAS covering a variety of human complex traits (**S3 Table**). We applied LD score regression [26] to stratify trait heritability by tissue and cell type, and identified a total of 226 significantly enriched annotation tracks for 34 traits after correcting for multiple testing (**S4-S7 Tables**). In general, GWAS with a large number of significant SNP-level associations showed stronger heritability enrichment in the predicted functional genome (**Figs 3A and 3B**). Tissue and cell tracks refined the resolution of heritability stratification and provided additional insights into the genetic basis of complex traits (**Figs 3C and 3D**).

The immune annotation track was significantly enriched for 7 immune diseases, namely celiac disease (CEL), Crohn’s disease (CD), ulcerative colitis (UC), primary biliary cirrhosis (PBC), rheumatoid arthritis (RA), systemic lupus erythematosus (SLE), and multiple sclerosis (MS). Using tracks for cell types, we identified several significant enrichments, including monocytes for CD (p=2.9e-11) and B cells for PBC (p=2.3e-6), RA (p=1.2e-5), and MS (p=2.2e-6). Inflammatory bowel diseases showed significant enrichment in the gastrointestinal (GI) annotation track (CD: p=1.4e-4; UC: p=5.6e-5). Another autoimmune disease with a well-established GI component, CEL, also showed nominal enrichment in the GI annotation track (p=3.7e-4).

Several brain annotation tracks were significantly enriched for associations of schizophrenia (SCZ), education years (EDU), and cognitive performance (IQ). Bipolar disorder (BIP), neuroticism (NEU), and chronotype (CHT) all showed nominally significant enrichment in the anterior caudate annotation track. Body mass index (BMI) and age at menarche (AAM) were significantly enriched in multiple brain annotation tracks. Compared to other brain regions, the substantia nigra annotation track showed weaker enrichment for these brain-based traits, which is consistent with its primary function of controlling movement.

Hundreds of height-associated loci have been identified in GWAS [27]. Such a highly polygenic genetic architecture is also reflected in our analysis. 59 of 66 tier-3 tissue and cell annotation tracks were significantly enriched for height associations, with breast myoepithelial cell (p=6.2e-14) and osteoblast (p=8.5e-14) being the most significant. Waist-hip ratio (WHR), birth weight (BW), and three blood pressure traits showed significant enrichment in the adipose annotation track. Overall, cardiovascular (CV) annotation tracks showed strong enrichment for blood pressure and coronary artery disease (CAD). Interestingly, the aorta annotation track is significantly enriched for pulse pressure (PP) but not systolic or diastolic blood pressure (SBP and DBP). CAD and 4 lipid traits, i.e. high and low density lipoprotein (HDL and LDL), total cholesterol (TC), and triglycerides (TG), shared a similar enrichment pattern in liver, adipose, and monocyte annotation tracks, which is consistent with the causal relationship among these traits [28].

Our results demonstrated that annotations with refined specificity could provide insights into disease etiology while broader annotations have greater statistical power. Age-related macular degeneration (AMD) was significantly enriched in broadly defined annotation tracks including immune, brain, CV, and GI, despite the non-significant enrichment results using tier-3 annotation tracks. Analyses based on all three tiers of annotations could systematically provide the most interpretable results for most traits. Importantly, we note that greater GWAS sample sizes will effectively increase statistical power in the enrichment analysis while leaving the overall enrichment pattern stable (**S1 Figure**). Therefore, many more suggestive enrichment results are likely to become significant as GWAS sample sizes grow. Finally, some traits, e.g. type-II diabetes (T2D) and age at natural menopause (AANM), showed strong enrichment in the general functional genome but not in specific tissues, suggesting that we may be able to gain a better understanding of these traits when annotation data for tissues or cell types more relevant to these traits are made available.

### Identify enrichment in immune-related DNA elements for neurodegenerative diseases

Next, we performed an integrative analysis of stage-I GWAS summary statistics from the International Genomics of Alzheimer’s Project [8] (IGAP; n=54,162) with GenoSkyline-Plus annotations (**Methods**). SNPs located in the broadly defined immune annotation track, which account for 24.4% of the variants in the IGAP data, could explain 98.7% of the LOAD heritability estimated using LD score regression (enrichment=4.0; p=1.5e-4). Somewhat surprisingly, the signal enrichment in DNA elements functional in immune cells was substantially stronger than the enrichment in brain and other tissue types (**Fig 4A**). To investigate if immune-related DNA elements are also enriched for associations of other neurodegenerative diseases, we analyzed a publicly accessible GWAS summary dataset for PD [29] (n=5,691; **Methods**). Again, the immune annotation track was the most significantly enriched annotation (enrichment=6.3; p=7.5e-6), followed by epithelium and CV (**Fig 4A**).

**Fig 4.**
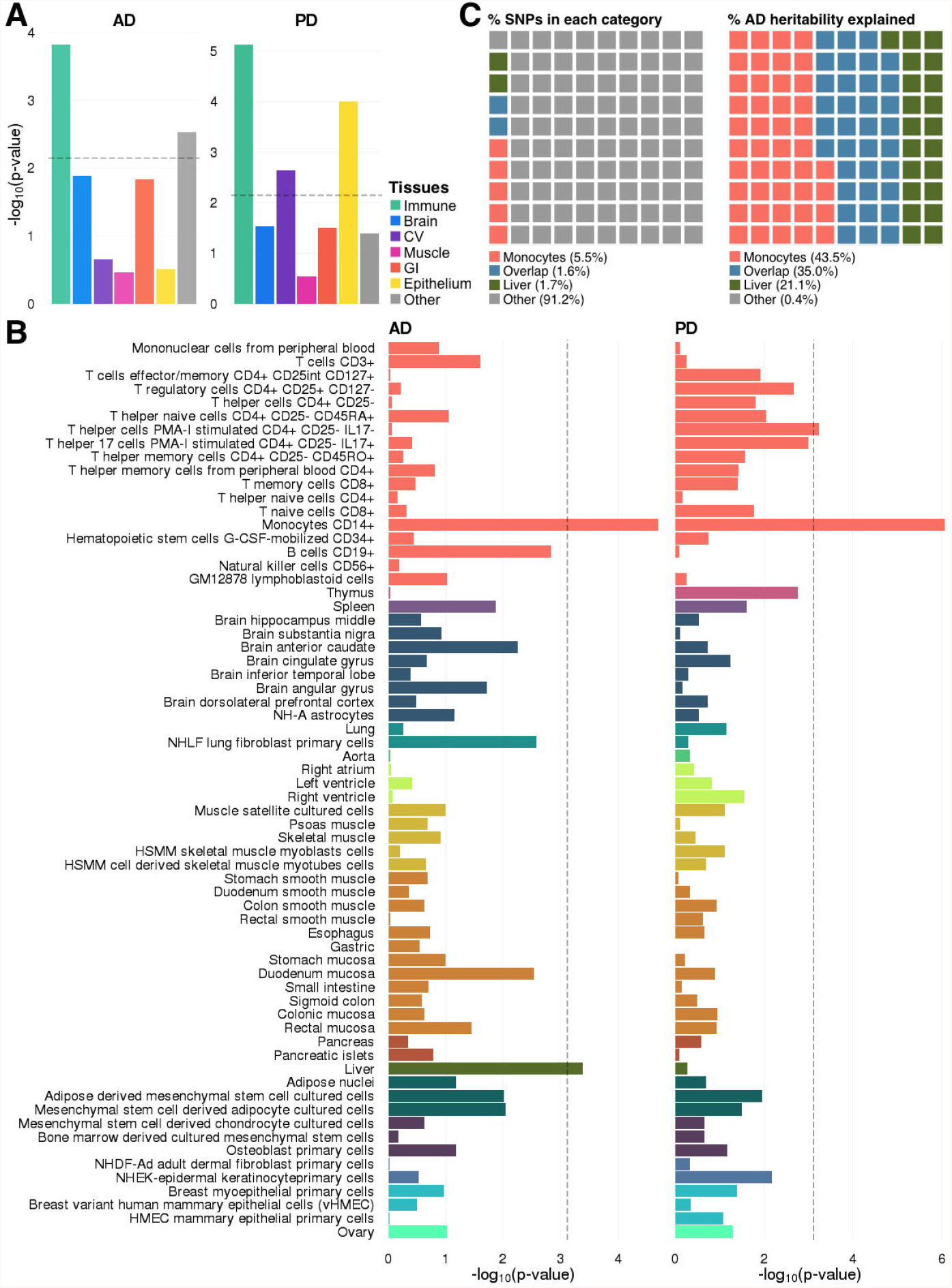
Tissue and cell type-specific enrichment for AD and PD. **(A)** Enrichment in 7 broadly defined tissue tracks. **(B)** Enrichment analysis using 66 GenoSkyline-Plus tissue and cell tracks. Dashed lines indicate Bonferroni-corrected significance cutoff. **(C)** Percentage of variants covered by each annotated category and percentage of heritability explained by variants in that category.

Analysis based on 66 tissue and cell tracks further refined the resolution of our enrichment study. Monocyte (enrichment=10.9; p=2.0e-5) and liver (enrichment=16.6; p=4.1e-4) annotation tracks were significantly enriched for LOAD associations (**Fig 4B**). In fact, the combined functional regions in monocyte and liver covered 8.8% of the SNPs in the IGAP data, but could account for 99.6% of the LOAD heritability currently captured in the IGAP stage-I GWAS (**Fig 4C**). In PD GWAS, signal enrichment in liver was absent, but monocyte-functional regions remained strongly enriched (enrichment=16.3; p=8.5e-7).

Our findings support the critical role of innate immunity in neurodegenerative diseases [10]. Significant enrichment for LOAD associations in liver-specific DNA elements also provides additional support for the possible involvement of cholesterol metabolism in LOAD etiology [30, 31]. LOAD signal enrichment in liver remained significant after removing the *APOE* region (chr19: 45,147,340-45,594,595; hg19) from the analysis (**S2 Figure**), suggesting a polygenic architecture in this pathway. Finally, some adaptive immune cells also showed enrichment for AD and PD associations. LOAD signal enrichment in the B cell annotation track was nominally significant, while multiple T cell annotation tracks were significantly enriched for PD associations. These results not only suggest the involvement of adaptive immunity in neurodegenerative diseases, but also hint at distinct mechanisms of such involvement between AD and PD. Finally, for comparison, we applied several other annotations including CADD [32], GWAVA [33], and EIGEN [34] to the LOAD GWAS data. GenoCanyon and GenoSkyline annotations for seven tissues were also included in the comparison. Our annotations outperformed these methods, showing stronger fold enrichment and more significant p-values (**S8 Table**).

### Identify shared genetic components between AD and PD

Our results showed strong enrichment for both AD and PD in the monocyte functional genome. Next, we investigate if the enrichment for both diseases is through shared or distinct genetic components. Recent studies have failed to identify statistically significant genome-wide pleiotropic effects between AD and PD [35]. We instead hypothesize that the same set of immune-related genetic components are involved in both diseases. Therefore, we aim to identify enrichment for pleiotropic effects in the genome localized to regions of monocyte functionality.

We first partitioned AD and PD heritability by chromosome. Chromosome-wide heritability showed moderate correlation between the two diseases (correlation=0.65; **Fig 5A**). When focusing on monocyte functional elements, chromosome-wide heritability showed high concordance between AD and PD (correlation=0.96; **Fig 5B**). Interestingly, such high concordance cannot be fully explained by chromosome size. In fact, the correlation between chromosome size and per-chromosome heritability estimates is 0.56 for AD and 0.59 for PD, both lower than the correlation between AD and PD’s per-chromosome heritability estimates, especially in the monocyte functional genome. The percentage of explained LOAD heritability on chromosome 19 is lower than previous estimation [36] due to removal of SNPs with large effects in the *APOE* region (**Methods**). Next, to quantify the shared genetics between AD and PD, we identified significant enrichment for pleiotropic effects in monocyte functional regions (enrichment=1.8; p=9.4e-4) using a window-based approach (**Methods**). To account for potential bias due to the moderate sample overlap between the two GWAS as well as other confounding factors, we applied a permutation-based testing approach (**Methods**). Enrichment for pleiotropic effects in the monocyte functional genome remained significant (p=4.6e-3). In addition, these results were robust with the choice of window size.

**Fig 5.**
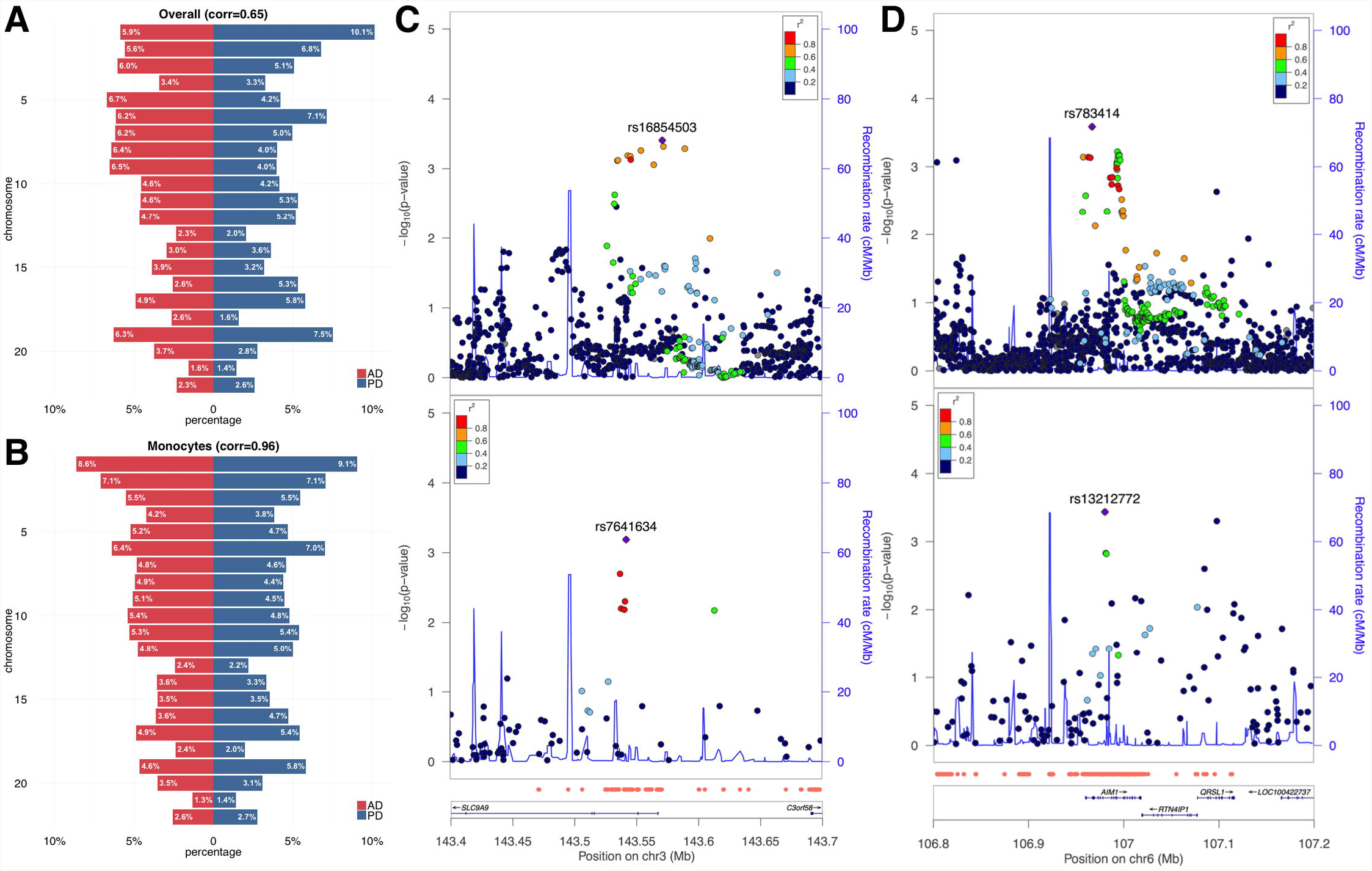
Identify genetic correlation between LOAD and PD. **(A)** estimated chromosome-by-chromosome heritability percentage for LOAD and PD. **(B)** chromosome-by-chromosome heritability in the monocyte functional genome. **(C-D)** Association peaks in pleiotropic loci *SLC9A9* and *AIM1*. The upper and the lower panels represent associations for LOAD and PD, respectively. Monocyte-specific functional regions are highlighted by red dots at the bottom of the figure above the gene annotations.

We identified 15 candidate loci for pleiotropic effects (**Methods**; **S9 Table**), among which signals at *SLC9A9* and *AIM1* are the clearest (**Figs 5C and 5D**). *SLC9A9*, whose encoded protein localizes to the late recycling endosomes and plays an important role in maintaining cation homeostasis (RefSeq, Mar 2012), is associated with multiple pharmacogenomic traits related to neurological diseases, including response to cholinesterase inhibitor in AD [37], response to interferon beta in MS [38], response to angiotensin II receptor blockade therapy [39], and multiple complex diseases including attention-deficit/hyperactivity disorder [40], autism [41], and non-alcoholic fatty liver [42]. Gene *AIM1* is associated with stroke [43], human longevity [44], and immune diseases including RA [45] and SLE [46].

A few candidate loci pointed to clear gene candidates but showed unclear or distinct peaks of association (**S3 Figure**). These include an inflammatory bowel disease risk gene *ANKRD33B* [47]. *PRUNE2* is a gene associated with response to amphetamine [48] and hippocampal atrophy which is a quantitative trait for AD [49]. *HBEGF* is associated with AD in *APOE* ε4-population [50] and involved in Aβ clearance [51]. *PROK2* is a gene involved in Aβ–induced neurotoxicity [52]. Additionally, the protein product of *AXIN1* negatively affects phosphorylation of tau protein [53]. Other gene candidates include *CCDC158*, *PRSS16*, and *ZNF615,* which are previously identified risk genes for PD, SCZ, and BIP, respectively [54-56]. Some other windows showed complex structures of linkage disequilibrium (LD) and contained large association peaks spanning a number of genes (**S4 Figure**), which include the region near PD risk gene *PRSS8* [54] and the *HLA* region. Interestingly, we also identified the surrounding region of *MAPT*, a gene that encodes the tau protein which is a critical component of both AD and PD pathologies [50, 54, 57, 58].

Pathway enrichment analysis for genes in 15 pleiotropic candidate loci identified significant enrichment in immune-related pathways staphylococcus aureus infection (KEGG:05150; p=1.9e-5) and systemic lupus erythematosus (KEGG:05322; p=3.7e-04; **Methods**). Both pathways remained significant after removing two *HLA* loci from our analysis.

### Reprioritize AD risk loci through integrative analysis of functional annotation

Finally, we reprioritize AD risk loci using monocyte and liver annotation tracks. We integrated IGAP stage-I summary statistics with GenoSkyline-Plus using genome-wide association prioritizer (GenoWAP [59]), and ranked all SNPs based on their GenoWAP posterior scores (**Methods**). Under a posterior cutoff of 0.95, we identified 8 loci that were not reported in the IGAP GWAS meta-analysis using monocyte annotation and 4 loci using the liver annotation track (**S10 Table**).

We then sought replication for SNPs with the highest posterior score at each of these loci using inferred IGAP stage-II z-scores (**Methods**). After removing shared SNPs between monocyte- and liver-based analyses, 10 SNPs remained in the analysis, 7 of which showed consistent effect directions between the discovery and the replication cohorts (**Fig 6A**). One SNP was successfully replicated in the inferred IGAP stage-II dataset, i.e. rs4456560 (p=0.013). SNP rs4456560 is located in *SCIMP* (**Fig 6B**), a gene that encodes a lipid tetraspanin-associated transmembrane adaptor protein that is expressed in antigen-presenting cells and localized in the immunological synapse [60].

**Fig 6.**
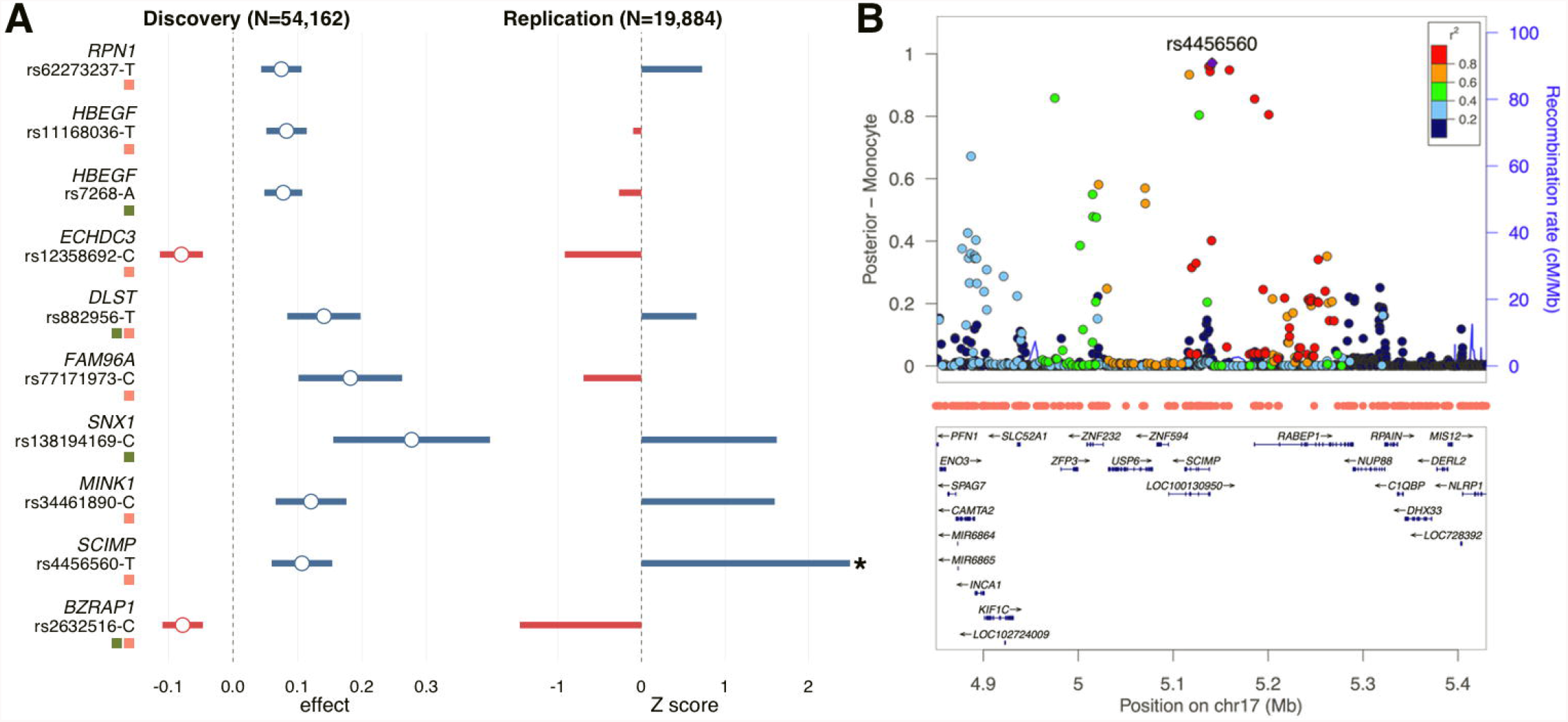
Reprioritize AD GWAS loci using functional annotations. **(A)** Effect size estimates for 10 SNPs of interest in the discovery and the replication cohort. Intervals in the discovery stage indicate 95% confidence. Asterisk indicates significant effects in the replication cohort. Red and green squares highlight loci identified using monocyte or liver annotation track, respectively. **(B)** The successfully replicated *SCIMP* locus. The vertical axis shows the GenoWAP posterior probability based on monocyte annotation track. Functional regions in monocyte are highlighted by red dots.

A moderate replication rate in the IGAP stage-II cohort was expected since we focused on loci that did not reach genome-wide significance in the IGAP meta-analysis and the IGAP stage-II cohort is relatively small (n=19,884) compared to the data in the discovery stage. Furthermore, data from IGAP stage-II cohort are not publicly available and we were limited to the inverse inference approach shown here. It is possible additional loci will replicate when IGAP stage-II summary or individual-level data are made available. However, all identified loci have been linked to AD or relevant phenotypes in the literature. *RPN1* was linked to AD through a network-based technique [61]. Association between *ECHDC3* and AD risk was established through a joint analysis of AD and lipid traits [62]. Association between *DLST* and AD has also been previously reported [63]. *BZRAP1* and *MINK1* were shown to be associated with cognitive function and blood metabolites, respectively [64, 65]. A pleiotropic effect candidate gene *HBEGF* showed up again in the SNP reprioritization analysis. Multiple genes in the sorting nexin family have been found to participate in *APP* metabolism and Aβ generation [66]. Association between *SNX1* and AD has also been previously identified using gene-based tests [67]. Finally, during the peer review process of this paper, three new genome-wide significant loci (i.e. *PFDN1/HBEGF*, *USP6NL/ECHDC3*, and *BZRAP1-AS1*) were reported in a trans-ethnic GWAS meta-analysis for AD [68], all of which were among our reprioritized list of risk loci. Further, the most significant SNPs at loci *PFDN1/HBEGF* (rs11168036, p=7.1e-9) and *BZRAP1-AS1* (rs2632516, p=4.4e-8) matched with our top reprioritized SNPs (**Fig 6A**).

## Discussion

Increasing evidence suggests that non-coding regulatory DNA elements may be the primary regions harboring risk variants in human complex diseases. In this work, we have substantially expanded our previously established GenoSkyline annotation by incorporating RNA-seq and DNA methylation into its framework, imputing incomplete epigenomic and transcriptomic annotation tracks, and extending it to more than 100 human tissue and cell types. With the help of integrative functional annotations, we identified strong enrichment for LOAD heritability in functional DNA elements related to innate immunity and liver tissue using hypothesis-free tissue-specific enrichment analysis. This enrichment was also found in immune-related DNA elements using PD data. Our analysis also clearly indicated that monocyte functional elements in particular appear to be highly relevant in explaining AD and PD heritability. Of note, we analyzed 45 complex diseases and traits in addition to AD and PD. The substantial enrichment for multiple psychiatric and neurological traits in the brain functional genome shows that the lack of brain enrichment in neurodegeneration is not due to poor quality of brain annotations. Further, the monocytes annotation track was the most significantly enriched for Crohn’s disease among the 45 GWAS, and was not ubiquitously enriched for a large number of traits. Consistent and biologically interpretable enrichment results on a large collection of complex traits demonstrate the effectiveness of our approach and increase the validity of novel findings.

It is worth noting that multiple studies have highlighted the role of myeloid cells in the genetic susceptibility of neurodegenerative diseases [11]. Several genes expressed in myeloid cells (e.g. ABCA7, CD33, and TREM2) have been identified in GWAS and sequencing-based association studies for AD [8, 69, 70]. Further, AD risk alleles identified in GWASs have been shown to enrich for cis-eQTLs in monocytes [9]. In addition, two recent papers identified enrichment for AD heritability in active genome regions in myeloid cells [71, 72], which suggested a polygenic genetic architecture for immune-related DNA elements in AD etiology and hinted at a large number of unidentified, immune-related genes for AD. Compared to the aforementioned work, our study utilizes a better set of tissue-specific genome annotations and explicitly accounts for the similarity between different cell types through a multiple regression model. One major limitation in our analysis is lack of data for other potentially AD-relevant cell types such as microglia. Whether our findings correctly reflected the direct involvement of peripheral immune cells in neurodegenerative diseases rather than the detection of epigenomic similarities between monocytes and microglia remains to be carefully investigated in the future.

Furthermore, we successfully identified enrichment for shared genetic components between AD and PD in the monocyte functional genome, which hints at a shared neuroinflammation pathway between these two neurodegenerative diseases. We note that several candidate loci with potential pleiotropic effects showed fairly marginal associations with AD and PD, which explains why they have been missed in traditional SNP-based association analysis. Importantly, SNPs in immune-related DNA elements explain a large proportion of AD and PD heritability in total. These results suggest that weak but pervasive associations related with immunity still remain unidentified. Further evaluations of these relationships using GWAS with larger sample sizes may provide insights into the shared biology of these neurodegenerative conditions.

Through multi-tier enrichment analyses on 45 GWAS, an in-depth case study of neurodegenerative diseases, and validation of known non-coding tissue-specific regulatory machinery, we have demonstrated the ability of GenoSkyline-Plus to provide unbiased, genome-wide insights into the genetic basis of human complex diseases. The analyzed GWAS represent a variety of human complex diseases and traits, highlighting the effectiveness of our method in different contexts and genetic architecture. However, while our non-coding validation study demonstrated that GenoSkyline-Plus annotations indeed captured tissue-specific activity in a variety of intergenic machinery, there is a need to develop a more statistically robust framework to identify new non-coding elements rather than validate existing ones. Our approach of identifying the functionally active proportion of all elements in aggregate is only able to identify tissue specificity while considering large groups of highly specific non-coding elements. The availability of over 100 different annotation tracks introduces many multiple-testing issues that should be addressed in the case of a statistically sound analysis for tissue-specificity. We have also demonstrated how GenoSkyline-Plus and its explanatory power improve with the addition of more data. Currently, functionality in 28% of exonic regions still remains to be identified. As the quantity and quality of high-throughput epigenomic data continue to grow, GenoSkyline-Plus has the potential to further evolve and provide even more comprehensive annotations of tissue-specific functionality in the human genome. We will update our annotations when data for new tissue and cell types from the Roadmap consortium become available. Finally, several recent papers have introduced novel models to integrate functional annotations in tissue-specific enrichment analysis [73, 74]. Many models that do not explicitly incorporate functional annotation information have also emerged in transcriptome-wide association studies and other closely-related applications in human genetics research [75-78]. Our annotations, in conjunction with rapidly advancing statistical techniques and steadily increasing sample sizes in genetics studies, may potentially benefit a variety of human genetics applications and promise a bright future for complex disease genetics research.

## Methods

### Annotation data preprocessing

Chromatin data were extracted from the Epigenomics Roadmap Project’s consolidated reference epigenomes database (http://egg2.wustl.edu/roadmap/). Specifically, ChIP-seq peak calls were collected for each epigenetic mark (H3k4me1, H3k4me3, H3k36me3, H3k27me3, H3k9me3, H3k27ac, H3k9ac, and DNase I Hypersensitivity) in each Roadmap consolidated epigenome where available. Peak calls imputed using ChromImpute [79] were used in place of missing data. Next, peak files were reduced to a per-nucleotide binary encoding of presence or absence of contiguous regions of strong ChIP-seq signal enrichment compared to input (Poisson p-value threshold of 0.01).

DNA methylation data were also collected from the Roadmap’s reference epigenomes database. CpG islands were identified in each sample using the CpG Islands Track of the UCSC Genome Browser (http://genome.ucsc.edu/), and unmethylated islands were those CpG islands with less than 0.5 fractionated methylation based on imputed methylation signal tracks in the Roadmap reference epigenomes database. Presence of an unmethylated CpG island was then encoded for each nucleotide as a binary variable. Finally, Roadmap’s RNA-seq data were dichotomized using an rpkm cutoff of 0.5 at 25-bp resolution and included in our annotations.

### GenoSkyline-Plus model

We adapt the existing framework established by Lu et al. to a broader set of genomic data [12]. Briefly, given a set of Annotations ***A*** and a binary indicator of genomic functionality ***Z***, the joint distribution of ***A*** along the genome is assumed to be a mixture of annotations at functional nucleotides and non-functional nucleotides. Assuming that each of the annotations in ***A*** is conditionally independent given ***Z*** conditionally independent given ***Z***, we factorize the conditional joint density of ***A*** given ***Z*** as:

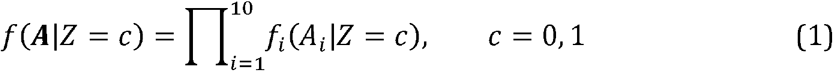

All annotations have been preprocessed into binary classifiers, and the marginal functional likelihood given each individual annotation can be modeled with a Bernoulli distribution

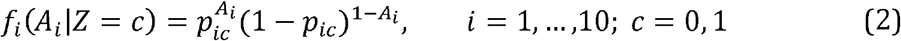

With an assumed prior probability π of functionality, the parameter *p_ic_* of each individual annotation can be estimated with the Expectation-Maximization (EM) algorithm. The posterior probability of functionality at a nucleotide, known as the GenoSkyline-Plus score, is then:

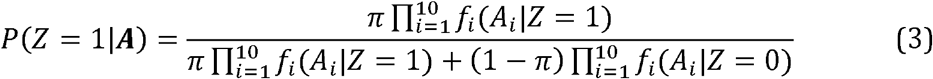

Giving us with 21 parameters for each annotation track:

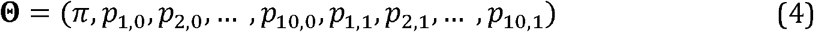

These parameters were estimated using the GWAS Catalog, downloaded from the NHGRI website (http://www.genome.gov/gwastudies/). 13,070 unique SNPs found to be significant in at least one published GWAS were expanded into 1kb bp intervals and formed a sampling covering 12,801,840 bp of the genome. This sampling method has been shown to be a robust representation of functional and non-functional regions along the genome [6]. Notably, other models have been recently developed to predict functional non-coding SNPs [34].

### Data for validating annotation quality

Quantile-normalized expression values were downloaded for all mature miRNAs profiled in Ludwig et al [17]. Due to inconsistent levels of miRNA specificity in the two donors in this study and to avoid diluting miRNA specificity, we used miRNA data from only body 1, which had a higher fraction of tissue specific miRNAs. TSI values were calculated as described in the study:

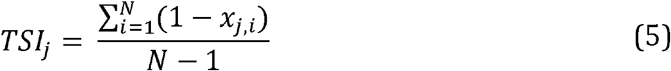

Where *N* is the total number of tissues measured, *x_j,i_* is the expression intensity of tissue *i* divided by the maximum expression across all tissues for miRNA *j*. We extract any miRNAs with a TSI score greater than the median value of 0.75 to produce a sufficiently large collection of miRNAs with expression highly specific to only a few tissues that we can then attempt re-identify using GenoSkyline-Plus. We next download genomic positions and identify the highest expressed tissue for each TSI-filtered miRNA. miRNA coordinates were extracted from miRbase (http://mirbase.org/) and mapped to hg19 using the UCSC liftover tool (http://genome.ucsc.edu/). lncRNA data was prepared similarly to miRNA. Expression data of 9,747 lncRNA transcripts based on GENCODE v3c annotation across 31 human tissues [19] (GEO accession: GSE34894) was downloaded. As above, the TSI of each lncRNA transcript was calculated, and transcripts with a TSI greater than 0.75 were labeled for genomic position and maximally expressed tissue.

Pre-defined enhancer differentially expressed cell facets [21] were downloaded from PrESSto database (http://enhancer.binf.ku.dk/presets/). Andersson et al. define their enhancer sets via bi-directional CAGE expression collected by the FANTOM consortium [80]. Cell facets were manually constructed using hierarchical FANTOM5 cell ontology term mappings to create mutually exclusive and broadly covered histological and functional annotations. Enhancers were considered differentially expressed in a facet using Kruskal-Wallis rank sum test and subsequent pair-wise post-hoc tests to identify enhancers with significantly differential expression between pairs of facets. Based on this method, an enhancer is considered differentially expressed in a facet if it is significantly differentially expressed compared to any other facet and has overall positive standard linear statistics.

For each of the three data validation sets, functional specificity is assessed by calculating the per-nucleotide functional proportion of all non-coding elements across a tissue. Functionality is defined by a Genoskyline-Plus score greater than 0.5 at that nucleotide. For Roadmap samples with multiple donors (e.g. skeletal muscle and rectal mucosa) we took the average GenoSkyline-Plus score at each nucleotide across the samples. For each set of non-coding elements we selected the top three tissues with the largest sample size that had matching annotations in Genoskyline-Plus. For example, we did not calculate scores for enhancers with maximal expressions in human testis because there is no corresponding Roadmap sample in which we would detect tissue-specific functionality.

To examine cell-specific functionality of the *IL17A* LCR in T-cell subsets, we extracted GenoSkyline-Plus scores for each nucleotide along the ∼200 kilobase region between the genes *PKHD1* and *MCM3* [23]. While scores for Th17 and Th1/Th2 subsets (i.e. ‘CD4+ CD25- IL17+ PMA-Ionomycin stimulated Th17 Primary Cells’ and ‘CD4+ CD25- IL17- PMA-Ionomycin stimulated MACS purified Th Primary Cells’; **S1 Table**) were extracted as-is, we took the average score of the two available CD4+ naïve T-cell subsets (i.e. ‘CD4 Naïve Primary Cells’ and ‘CD4+ CD25- CD45RA+ Naïve Primary Cells’). We identified the analogous human regions of previously identified functional murine CNS regions [25] by taking the top 20 most conserved intergenic sites between mouse and human in the LCR region using the VISTA browser (http://pipeline.lbl.gov/cgi-bin/gateway2). GenoSkyline-Plus scores in the 20 CNS sites and their 200-bp flanking regions were compared across different cell types.

### GWAS data details

Summary statistics for 45 GWAS are publicly accessible. Details for these studies are summarized in **S3 Table**. IGAP is a large two-stage study based upon genome-wide association studies (GWAS) on individuals of European ancestry. In stage-I, IGAP used genotyped and imputed data on 7,055,881 SNPs to meta-analyze four previously-published GWAS datasets consisting of 17,008 Alzheimer's disease cases and 37,154 controls (The European Alzheimer's disease Initiative – EADI, the Alzheimer Disease Genetics Consortium – ADGC, The Cohorts for Heart and Aging Research in Genomic Epidemiology consortium – CHARGE, and The Genetic and Environmental Risk in AD consortium – GERAD). In stage-II, 11,632 SNPs were genotyped and tested for association in an independent set of 8,572 AD cases and 11,312 controls. Finally, a meta-analysis was performed combining results from stages I and II. IGAP stage-I GWAS summary data is publicly accessible from IGAP consortium website (http://web.pasteurlille.fr/en/recherche/u744/igap/igap_download.php). GWAS summary statistics for PD was acquired from dbGap (accession: pha002868.1). Details for AD and PD studies have been previously reported [8, 29].

### Stratify heritability by tissue and cell type

Heritability stratification and enrichment analyses were performed using LD score regression implemented in the LDSC software (https://github.com/bulik/ldsc/). Annotation-stratified LD scores were estimated using dichotomized annotations, 1000 Genomes (1KG) samples with European ancestry [81], and a default 1-centiMorgan window. Enrichment was defined as the ratio between the percentage of heritability explained by variants in each annotated category and the percentage of variants covered by that category.

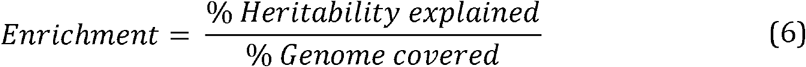

A resampling-based approach was used to assess standard error estimates [26]. Three tiers of annotations of different resolutions were used in enrichment analyses:

1. Generally functional genome predicted by GenoCanyon annotation smoothed along 10-kb windows.
2. Seven unique tissue and cell type clusters (i.e. immune, brain, CV, muscle, GI, epithelium, and other), representing common, physiologically related organ systems. Each category is defined as the union of functional regions in related tissue and cell types (**S2 Table**).
3. GenoSkyline-Plus annotations for 66 selected tissue and cell types (**S2 Table**).

The smoothing strategy for GenoCanyon improves its ability to identify general functionality in the human genome [59]. GenoSkyline-Plus and smoothed GenoCanyon annotations were dichotomized using a cutoff of 0.5. Such dichotomization is robust to the cutoff choice due to the bimodal nature of annotation scores [6]. We selected 66 annotation tracks in the tier-3 analysis by removing all the fetal and embryonic cells, and taking the union of different Roadmap epigenomes for the same cell type (**S2 Table**). The 53 baseline annotations of LD score regression were always included in the model across all analyses as suggested in the LDSC user manual. Smoothed GenoCanyon annotation track was also included in tier-2 and tier-3 analyses to account for unobserved tissue and cell types. Of note, the proposed multiple regression model explicitly takes the overlapped functional regions across biologically-related cell types into account. Further, the linear mixed-effects model in LDSC does not assume linkage equilibrium, and therefore LD will most likely not introduce bias into heritability estimation and enrichment calculation. We removed the MHC region from our analysis due to its unique LD patterns.

A slightly different strategy was adopted when comparing the performance of different computation annotation tools. To make fair comparison, we dichotomized all annotation tracks using each score’s top 90% quantile calculated from SNPs with minor allele count greater than five in 1000 Genomes samples with European ancestry. We then followed the suggested protocol of LDSC and kept baseline annotations in the model while adding each annotation track one at a time.

### Pleiotropy analysis

We calculated chromosome-by-chromosome heritability percentage through summing up and normalizing per-SNP heritability estimated using LD score regression and tier-3 annotation tracks. Of note, since only GWAS summary statistics were used as the input, popular heritability estimation tools such as GCTA [82] could not be applied. The sums over complete chromosomes are compared with the sums over monocyte functional regions only. Notably, LDSC is conceptually different from some other tools (e.g. GCTA [82]) in its estimation of trait heritability. GCTA estimates the proportion of phenotypic variability that can be explained by SNPs in the GWAS dataset while LDSC aims to estimate the proportion of phenotypic variability explained by all the SNPs in samples from the 1KG Project. In practice, LDSC only uses HAPMAP SNPs to fit the LD score regression model and assumes that HAPMAP SNPs are sufficient for tagging all 1KG SNPs through LD [26]. Additionally, LDSC applies a few stringent SNP filtering steps for quality control reasons, e.g. removing SNPs with very large effect sizes (i.e χ^2^ > 80), which leads to the removal of some SNPs in the *APOE* region in our analysis.

To evaluate enrichment of pleiotropic sites in the monocyte functional genome, we partition the genome into windows with length of 1M bases. Sex chromosomes and windows without SNPs are removed in our datasets. For each disease (i.e. AD and PD), we label a window 1 if the following criteria are met.

1. There is at least one SNP with p-value < 1e-3 in the window.
2. Among SNPs that meet condition 1, at least one is located in the monocyte-specific functional genome.

Otherwise, the window is labeled 0. This labeling results in two binary vectors, one for each disease. A window marked as 1 for both AD and PD is a window of interest that suggests a possible association in monocytes-related DNA for both diseases in that region. We use a hypergeometric test to assess if such a pattern of local association appears more often than by chance. Windows marked as 1 for both diseases are subsequently curated to identify the association peaks that potentially have pleiotropic effects for AD and PD.

There is a moderate overlap of control samples between IGAP AD GWAS and the PD GWAS (KORA controls, N∼480). To account for the bias introduced by sample overlap and other confounding factors, we designed a permutation-based approach. In each permutation step, we shuffle the annotation status while keeping the total proportion of annotated regions, and then pick out windows that meet condition 2. We calculate the p-value through comparing the observed number of windows that meet conditions 1 and 2 for both diseases with the empirical distribution acquired in permutations.

Of note, we also applied this approach using a window size of 500K bases. Results in all related tests remained similar.

### GWAS loci reprioritization

We briefly describe the SNP reprioritization approach implemented in the GenoWAP software available on our server (http://genocanyon.med.yale.edu/GenoSkyline). First, we identify three disjoint cases for SNPs in a given GWAS dataset.

1. The SNP is in a genomic region that is functional for the given phenotype and tissue (*Z_D_* = 1, *Z_T_* = 1).
2. The SNP is in a genomic region that is functional in the given tissue, but that region has no functionality for the phenotype (*Z_D_* = 0, *Z_T_* = 1).
3. The SNP is in a genomic region that is not functional in the given tissue (*Z_T_* = 0).

A useful metric for prioritizing SNPs is the conditional probability that the SNP is classified under case-1 given its p-value in the GWAS study, i.e. *P*(*Z_D_* = 1, *Z_T_* = 1 | *p*). We can denote this probability using Bayes formula as follows:

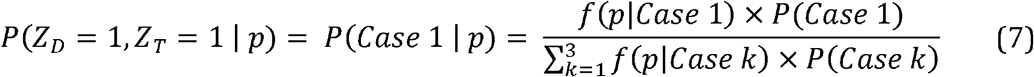

First, *P*(*Case* 3) = 1 − *P*(*Z_T_* = 1) can be directly identified using GenoSkyline-Plus scores. We partition all the SNPs into two subgroups based on a mean GenoSkyline-Plus score threshold of 0.1. Notably, these probabilities are not sensitive to changing threshold [6]. In this way, we can directly estimate *f*(*p*|*Case* 3)=*f*(*p*|*Z_T_* = 0) by applying a histogram approach on the SNP subgroup with low GenoSkyline-Plus scores.

Next, we assume that SNPs that are functional in a tissue but not relevant to the phenotype will have the same p-value distribution to all other SNPs that are not relevant to the phenotype, which in turn behave similarly to SNPs that are not functional at all. We have previously demonstrated that this assumption is backed by empirical evidence [6]. More formally, this relationship is denoted as follows:

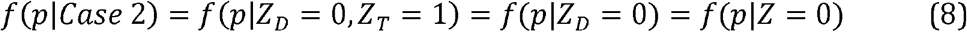

We estimate the distribution *f*(*p*|*Z* = 0) by using a similar approach to estimating *f*(*p*|*Z_T_* = 0, but partitioning SNPs using the general functionality GenoCanyon score instead of tissue-specific GenoSkyline-Plus score.

Finally, all remaining terms in formula 6 can be estimated using the EM algorithm. The p-value distribution of the subset of SNPs located in tissue-specific functional regions (i.e. *Z_T_* = 1) is the following mixture:

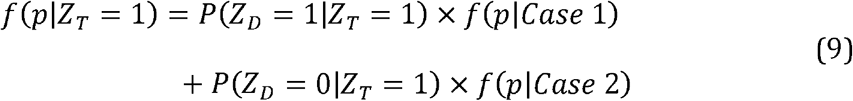

Density function *f*(*p*|*Case* 2) has been estimated in formula (8) and *f*(*p*|*Case* 2) is assumed to follow a beta distribution, which guarantees a closed-form expression in the EM algorithm.

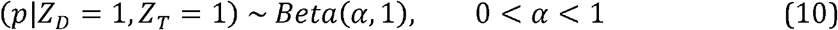

Notably, the *APOE* region was removed in the SNP reprioritization analysis for LOAD.

### Inverse inference of IGAP stage-II z-scores

Summary statistics from both IGAP stage-I GWAS and stage-I + II meta-analysis are publicly available (http://web.pasteur-lille.fr/en/recherche/u744/igap/igap_download.php). We inferred z-scores from IGAP stage-II replication cohort using the following formula.

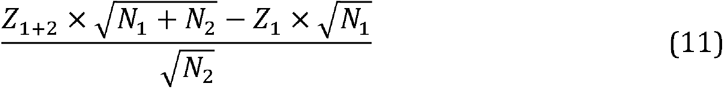

In this formula, *Z*_1_ and *Z*_1 + *z*_ indicate z-scores from the stage-I GWAS and the combined meta-analysis, respectivel *N_i_* indicates the sample size from the i^th^ stage. This formula was derived from the sample size based meta-analysis model, an approach known to be asymptotically equivalent to inverse variance based meta-analysis [83].

### Data accessibility and other bioinformatics tools

GenoSkyline-Plus annotation tracks, tiers 1-3 LD score files, and scripts for generating GenoSkyline-Plus scores are freely available on the GenoSkyline server (http://genocanyon.med.yale.edu/GenoSkyline). All annotation tracks can be visualized using UCSC genome browser. Web server g:Profiler was used to perform pathway enrichment analysis [84]. The g:SCS threshold implemented in g:Profiler was applied to account for multiple testing. Locus plots were generated using LocusZoom [85]. Gene plots were generated using R package “Gviz”.

## Acknowledgements

We thank the International Genomics of Alzheimer's Project (IGAP) for providing summary results data for these analyses. The investigators within IGAP contributed to the design and implementation of IGAP and/or provided data but did not participate in analysis or writing of this report. IGAP was made possible by the generous participation of the control subjects, the patients, and their families. We are also grateful for all the consortia and investigators that provided publicly accessible GWAS summary statistics.

## Supporting information captions

**S1 Figure. Enrichment analysis for three schizophrenia studies.** Overall enrichment pattern for schizophrenia remains stable as sample size increases. Sample sizes for the three studies shown below are 21,856, 32,143, and 82,315, respectively.

**S2 Figure. Enrichment analysis for LOAD after removing the APOE region.**

**S3 Figure. Nine pleiotropic loci for LOAD and PD.** For each locus, the upper and lower panels show associations for LOAD and PD, respectively. Monocyte functional regions are marked by red dots above gene names.

**S4 Figure. Four pleiotropic loci that span a large number of candidate genes.** For each locus, the upper and lower panels show associations for LOAD and PD, respectively. Monocyte functional regions are marked by red dots above gene names.

**S1 Table. List of 127 GenoSkyline-Plus annotation tracks.** Epigenome ID and cell type list are acquired from Epigenomics Roadmap Project and ENCODE.

**S2 Table. Curated annotation tracks used in tissue-specific enrichment analysis.**

**S3 Table. Details of 45 genome-wide association studies.**

**S4 Table. P-values for tissue-specific enrichment analysis (tiers 1 and 2).**

**S5 Table. Fold change for tissue-specific enrichment analysis (tiers 1 and 2).**

**S6 Table. P-values for tissue-specific enrichment analysis (tier 3).**

**S7 Table. Fold change for tissue-specific enrichment analysis (tier 3).**

**S8 Table. AD enrichment in other computational annotations.**

**S9 Table. Candidate loci for pleiotropic effect between AD and PD.**

**S10 Table. Top loci based on GenoWAP posterior scores.** Subscripts indicate stage-I discovery cohort, stage-II replication cohort, or stage-I + II meta-analysis. Each locus may contain multiple genes. SNP rs143560707 was not genotyped in the IGAP stage-II cohort.

